# Origin and evolution of the Zinc Finger Antiviral Protein

**DOI:** 10.1101/2020.12.04.412510

**Authors:** Daniel Gonçalves-Carneiro, Matthew A. Takata, Heley Ong, Amanda Shilton, Paul D. Bieniasz

## Abstract

The human zinc finger antiviral protein (ZAP) recognizes RNA by binding to CpG dinucleotides. Mammalian transcriptomes are CpG-poor, and ZAP may have evolved to exploit this feature to specifically target non-self viral RNA. Phylogenetic analyses reveal that *ZAP* and its paralogue *PARP12* share an ancestral gene that arose prior to extensive eukaryote divergence, and the *ZAP* lineage diverged from the *PARP12* lineage in tetrapods. Notably, The CpG content of modern eukaryote genomes varies widely, and *ZAP*-like genes arose subsequent to the emergence of CpG-suppression in vertebrates. Human PARP12 exhibited no antiviral activity against wild type and CpG-enriched HIV-1, but ZAP proteins from several tetrapods had antiviral activity when expressed human cells. In some cases, ZAP antiviral activity required a TRIM25 protein from the same or a related species, suggesting functional co-evolution of these genes. Indeed, a hypervariable sequence in the N-terminal domain of ZAP contributed to species-specific TRIM25 dependence in antiviral activity assays. Crosslinking immunoprecipitation coupled with RNA sequencing revealed that ZAP proteins from human, mouse, bat and alligator exhibit a high degree of CpG-specificity, while some avian ZAP proteins appear more promiscuous. Together, these data suggest that the CpG-rich RNA directed antiviral activity of ZAP-related proteins arose in tetrapods, subsequent to the onset of CpG suppression in certain eukaryote lineages, with subsequent species-specific adaptation of cofactor requirements and RNA target specificity.

**Author Summary:** To control viral infections, cells have evolved a variety of mechanisms that detect, modify and sometimes eliminate viral components. One of such mechanism is the Zinc Finger Antiviral Protein (ZAP) which binds RNA sequences that are rich in elements composed of a cytosine followed by a guanine. Selection of viral RNA can only be achieved because such elements are sparse in RNAs encoded by human genes. Here, we traced the molecular evolution of ZAP. We found that ZAP and a closely related gene, PARP12, originated from the same ancestral gene that existed in a predecessor of vertebrates and invertebrates. We found that ZAP proteins from mammals, birds and reptiles have antiviral activity but only in the presence of a co-factor, TRIM25, from the same species. ZAP proteins from birds were particularly interesting since they demonstrated a broader antiviral activity, primarily driven by relaxed requirement for cytosine-guanine. Our findings suggest that viruses that infect birds – which are important vectors for human diseases – are under differential selective pressures and this property may influence the outcome of interspecies transmission.

## Introduction

Organisms have evolved numerous mechanisms to detect and control viral infections. For example, pattern recognition receptors (PRRs), including RIG-I-like receptors and Toll-like receptors, can recognize RNA or DNA structures that are uniquely present or inappropriately localized in virus-infected cells (1). Recognition by PRRs triggers a signalling cascade that culminates in the increased transcription of many so-called interferon-stimulated genes (ISGs), some of which encode effectors with direct antiviral properties (2). PRRs, signalling molecules and direct antiviral effectors often exhibit species-dependent sequence and functional divergence, as a natural consequence of extreme reciprocal selective pressures placed on hosts and the viruses that colonize them.

The zinc finger antiviral protein (ZAP) is unusual in that it combines features of a nucleic acid PRR and a direct antiviral effector. ZAP was initially found to inhibit the replication of broad range of unrelated viruses, and to act by destabilizing viral RNA (3–5). Human ZAP is composed of three structural domains: an N-terminal RNA-binding domain that has four CCCH-type zinc fingers, a central domain that has an additional CCCH-type zinc finger plus two WWE domains, and a C-terminal poly(ADP-ribose)-polymerase (PARP)-like domain. ZAP requires certain cofactors for its antiviral activity, including TRIM25, a E3 ubiquitin ligase that interacts with ZAP via its SPRY domain (6,7). However, the precise role of TRIM25 in ZAP function is unknown. Several helicases and ribonucleases, including the putative endonuclease KHNYN, have also been reported to be required for ZAP activity (8,9).

We recently showed that ZAP targets particular RNA elements based on their dinucleotide composition (10). Specifically, ZAP binds directly to RNA elements that contain CpG dinucleotides, and RNAs that contain CpG-rich sequences exhibit cytoplasmic depletion in the presence of ZAP. X-ray crystal structures of the N-terminal RNA binding domain in a complex with an RNA target have revealed that the specificity of ZAP for CpG dinucleotides is conferred by a binding pocket positioned within a larger RNA binding domain (11,12). This pocket can only accommodate CpG dinucleotides, and its integrity is required for specific binding to CpG-rich RNA. Accordingly, the CpG content of viral genomes predicts their sensitivity/resistance to ZAP (10).

Human ZAP has minimal effects on the host transcriptome, presumably because CpG dinucleotides have been largely purged from the human genome, rendering human mRNAs largely ZAP-resistant. The purging of CpG dinucleotides (or ‘CpG-suppression’) from host genomes has occurred through DNA methylation followed by spontaneous deamination at CpG dinucleotides over millions of years and to varying degrees in different lineages (13). A commonly accepted theory that would enable this phenomenon argues that the emergence of DNA methyltransferases – enzymes that catalyse the conversion of cytosines in a 5’-cytosine-guanine-3’ (CpG) context to 5-methyl-cytosine – led to increased levels of methylated CpG. While spontaneous deamination of the methylated cytosine generates thymine, an authentic DNA base, deamination of an unmethylated cytosine generates deoxyuridine, that would be repaired. Thus, many DNA genomes have become purged of CpG dinucleotides and enriched in TpG dinucleotides (14).

Remarkably, the genomes of many RNA viruses are CpG-poor, and to a large extent CpG suppression in host is mirrored by CpG suppression in the viruses that colonize them. This feature is true even for RNA viruses whose genome composition could not have been be shaped by DNA methylation/deamination. The selective pressures that led to CpG suppression in viral genomes remain unknown, but long-term adaptation to hosts in which a selective pressure was imposed by ZAP is a likely possibility.

Here we investigated the origins of ZAP and its CpG-specific antiviral activity. A paralogue of *ZAP*, *PARP12*, exists that also lacks poly(ADP-ribose) polymerase activity, but catalyses the addition of mono(ADP-ribose) to proteins (15). Like ZAP, PARP12 also contains five CCCH-type zinc fingers, two WWE domains and has been reported to exhibit antiviral activity against several RNA viruses including vesicular stomatitis virus (VSV), ECMV and Dengue virus (16–18). Since *ZAP* and *PARP12* share domain and sequence homology, it is likely that these antiviral genes have share the same ancestral gene. However, when this gene duplication occurred and when specific CpG-binding affinity of ZAP-like proteins emerged remains unknown. We examined the genomes of vertebrates and invertebrates for gene products with sequence and domain homology to ZAP/PARP12. Phylogenetic analysis of *ZAP/PARP12*-related sequences showed that these genes arose early in the divergence of vertebrates and have their origin in an ancestral gene whose decedents are present in some modern invertebrates, such as cnidarians, but absent in others, such as arthropods. Each of the ZAP -related proteins from tetrapods that were examined showed antiviral activity against CpG-enriched HIV-1 when overexpressed in human cells. Notably, in some cases co-expression of a cognate TRIM25 protein in human cells was essential for the activity of ZAP proteins from non-mammalian species (e.g., chicken and alligator) suggesting that these proteins have co-evolved. Finally, we found that while CpG targeting specificity is common among ZAP proteins of human, mouse, bats and alligator origin, ZAP proteins of avian origin were less specific for CpG enriched targets and inhibited the replication of both wild type and mutant HIV-1. These data suggest lineage-specific evolution of ZAP and its cofactors with distinct non-self RNA targeting abilities that could influence the host range of viruses with zoonotic potential.

## RESULTS

### The dinucleotide composition of mRNAs and the origin of ZAP-like genes

To understand the evolutionary origins of ZAP-mediated, CpG-rich RNA targeted, non-self RNA recognition, we first mined vertebrate and invertebrate genomes for gene products with sequence homology to human ZAP. In the human genome, a *ZAP* homolog, *PARP12,* that shares several features with ZAP including five CCCH-type zinc fingers, two WWE domains and a C-terminal PARP domain (**Fig 1A,S1 Fig A and B**), is found at a proximal location on chromosome 7 (**Fig 1B**), suggesting that *ZAP* and *PARP12* are paralogs that arose from a common ancestor. A related gene termed ‘ZAP-like’ is present on human chromosome 7 but contains sequences corresponding only to the ZAP N-terminal RNA binding domain (**Fig 1A,S1 Fig A and B**). For searches of other genomes we considered sequences that contained five CCCH-type zinc fingers and a WWE domain in a ZAP/PARP12-like configuration (**Fig 1A**). Most of the sequences revealed by blast searches also encoded PARP-like domains, in common with ZAP and PARP12, but the PARP domain was lacking in ZAP/PARP12 sequences from a few species. The topology of a phylogenetic tree constructed using ZAP/PARP12 protein sequences revealed two separate lineages in vertebrates and invertebrates (**Fig 1C**) suggesting that a gene that is ancestral to modern *ZAP/PARP12* originated prior to the divergence of cnidarians from other animals, approximately 650 million years ago. While *ZAP/PARP12*-related sequences were found in several invertebrate genomes, the *Arthropoda* phylum, which includes insects and nematodes, appeared to lack *ZAP/PARP12* homologs, suggesting loss of *ZAP/PARP12* in this lineage.

**Fig 1.**
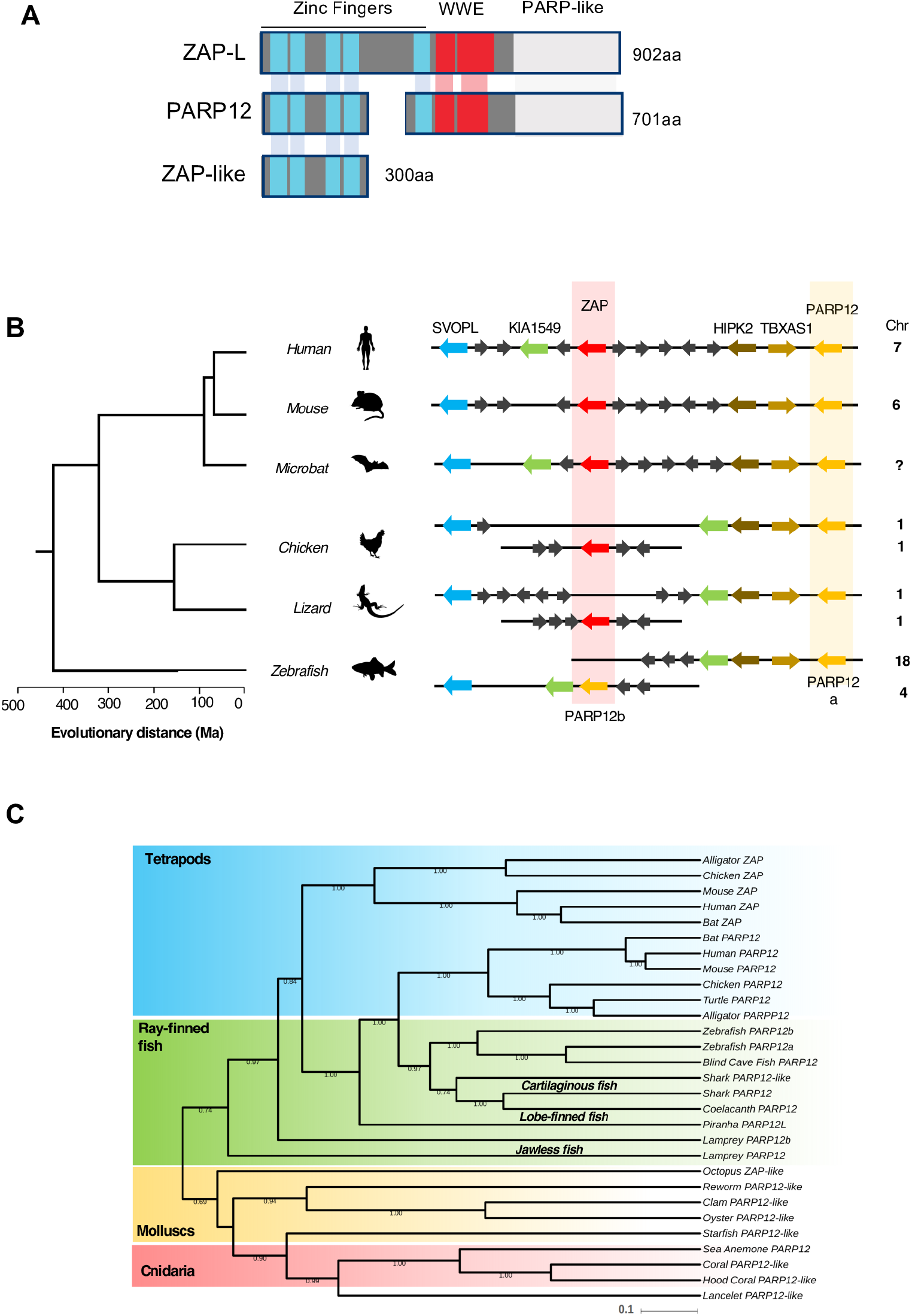
ZAP and PARP12 share a common ancestor. (A) Schematic representation of the domain organization of human ZAP and its paralogues PARP12 and ZAP-like. (B) Diagrams of the organization of PARP12 (in yellow) and ZAP (in red) loci in vertebrate genomes generated using multiple genome browsers (NCBI, Ensembl, Genomicus). Evolutionary distance of the indicated species (presented in million years, Ma) is based on (34). Chr, chromosome number. (C) Maximum likelihood tree (midpoint rooted) depicting phylogenetic relationships among *PARP12-* and *ZAP*-related genes in vertebrate and invertebrate species, generated with BEAST and 1000 bootstrap replicates. Shaded areas loosely cluster sequences from related species in the superclass Tetrapoda (in blue), clade Actinopterygii (Ray-finned fish, in green), the phylum Mollusca (molluscs, in yellow) and the phylum Cnidaria (in red).

Invertebrate genomes typically contained only one copy of a *ZAP/PARP12*-related gene, while vertebrates had two (**Fig 1B, C**). One set of *ZAP/PARP12* related sequences were more closely related to PARP12, and appeared to be represented once in tetrapod genomes and twice in the genomes of several fish genomes (**Fig 1 B and C,S1 Fig C**), suggesting at least one duplication event that was preserved in fish lineages. A clearly distinguishable set of *ZAP/PARP12* genes emerged exclusively in tetrapods, and this linage contained the human *ZAP* gene that has been demonstrated to exhibit antiviral activity against CpG-rich viruses (**Fig 1C**). Within tetrapods, the *ZAP*-related genes were more diverse in sequence than the PARP12-related sequences (**Fig 1C,S1 Fig C**). Overall these data suggest that ZAP originated in tetrapods from the duplication of an ancestral *PARP12* gene, and that *ZAP* may have been under stronger diversifying selective pressure than *PARP12* (19).

Consistent with the aforementioned scenario, *ZAP* is located at about 900kb upstream of *PARP12* in mammalian genomes, while *ZAP* and *PARP12* are further apart but on the same chromosome in reptilian and avian genomes (**Fig 1B**). In the zebrafish genome, duplicated *PARP12a* and *PARP12b* genes are located on two different chromosomes (18 and 4, respectively, **Fig 1B**). The *PARP12a* locus resembles the *PARP12* locus in birds and reptile while the *PARP12b* locus resembles that of *ZAP* in mammalian genomes (**Fig 1B**). Thus, even though CpG-suppression is commonly observed among vertebrates, the genes most closely related to human *ZAP* are found only in tetrapods (mammals, birds and reptiles).

### Human PARP12 does not recognize or inhibit expression from CpG-rich RNA

Since ZAP and PARP12 share an ancestral origin and are structurally related (**Fig 1**), we inquired if PARP12 also had antiviral activity against CpG-enriched HIV-1. To assess this, we measured the infectious virus yield and viral protein expression in HEK293T ZAP^−/−^ cells co-transfected with wild-type HIV-1 (HIV-1_WT_) or CpG-enriched mutant (HIV-1_CG_) proviral plasmids and increasing amounts of plasmids expressing either human ZAP-L or human PARP12. Increasing levels of ZAP-L reduced HIV-1_CG_ Env levels and the yield of infectious HIV-1_CG_ but did not affect HIV-1_WT_, as expected, but PARP12 had no effect on the yield or protein expression by either virus (**Fig 2A and B**). To test effects on other CpG-enriched or CpG-suppressed virus derived sequences, we used a luciferase-based reporter system in which ZAP or PARP12 expression plasmids were cotransfected with a plasmid encoding the firefly luciferase gene and a 3’ UTR containing vesicular stomatitis virus (VSV) or influenza A virus (IAV) derived sequences (**Fig 2C**). Plasmids that encoded CpG-enriched derivatives of these sequences were included in the experiment (10). Overexpression of ZAP specifically reduced the expression of the reporter gene with CpG-enriched 3’UTR sequences; however, PARP12 had no effect (**Fig 2C**). Thus, while they are structurally related, PARP12 does not share the same antiviral activity as ZAP. Differences in antiviral activity or specificity may be attributable to variation in RNA-binding activity, as ZAP:RNA adducts were easily detected in crosslinking immunoprecipitation assays, while PARP12:RNA adducts were undetectable under the same experimental conditions (**Fig 2D**).

**Fig 2.**
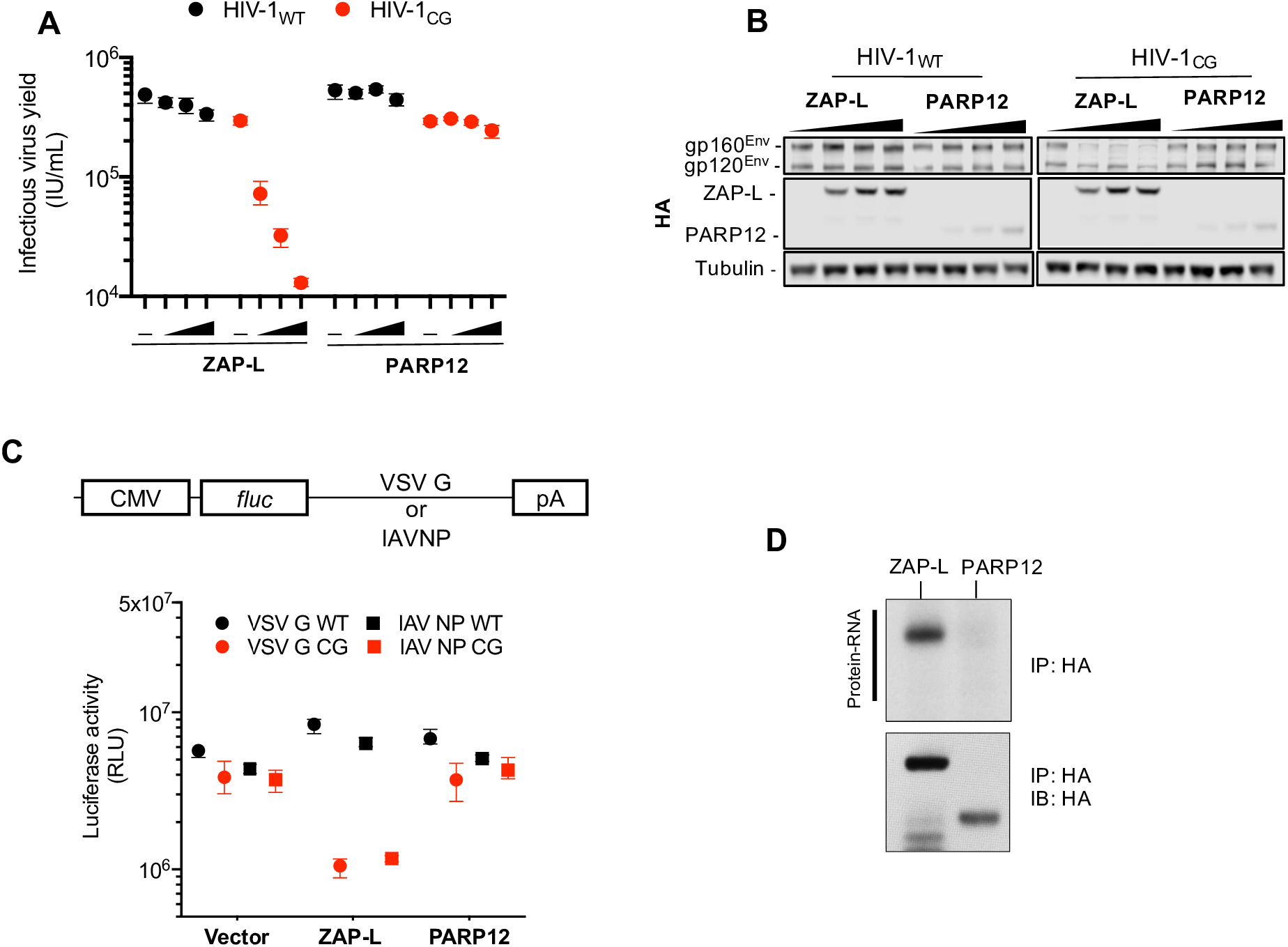
Human PARP12 does not restrict CpG-rich viruses. (A-B) HEK29T ZAP^−/−^ cells were transfected with HIV-1_WT_ or HIV-1_CG_ plasmids and increasing amounts (0ng, 75ng, 145ng, or 290ng) of a plasmid encoding human ZAP-L-HA or PARP12-HA. Supernatant was harvested after 48h and the infectious virus yield was determined using MT4-R5-GFP target cells (A). Whole cell lysates were analysed by western blotting (B). IU, infectious units. (C) HEK293T ZAP^−/−^ cells were transfected with plasmids encoding a luciferase reporter gene that contained VSV-G wildtype (WT) or CG-enriched (VSV-G CG) sequences, or IAV-NP WT or CG-enriched sequences as 3’ UTRs, together with plasmids expressing ZAP-L, PARP12 or an empty plasmid (vector). RLU, relative light units. (D) HEK293T ZAP^−/−^ cells were transfected with plasmids expressing ZAP-L-3xHA or PARP12-3xHA and 24h later culture medium was supplemented with 100μM 4SU. After overnight incubation cells were irradiated with UV light, and ZAP-L/PARP12 were immunoprecipitated. RNA bound to each protein was radiolabeled and protein-RNA adducts were resolved by SDS-PAGE, transferred to a nitrocellulose membrane and exposed to autoradiographic film. In parallel, a western blot of immunoprecipitated proteins was done using anti-HA antibody. IP, immunoprecipitation. IB, immunoblot.

These data suggest the that CpG-specific antiviral activity emerged concurrently with or after the emergence of the ZAP-like genes from a ZAP/PARP12 ancestral progenitor. To assess whether the emergence of ZAP-like proteins was potentially enabled by the prior presence CpG suppression in eukaryotic genomes, we generated an *in silico* library of open reading frames (ORFs) and 3’UTRs from organisms across the tree of life, including: vertebrates, nematodes, insects, molluscs and plants. We first calculated the frequency distribution (observed/expected) for each of the 16 possible dinucleotides in ORFs (**Fig 3A**). Notably, dinucleotide frequencies among distantly related species were similar, with the exception of the dinucleotide CpG. Specifically, ORF sequences from arthropods and nematodes had CpG frequencies close to that expected based on their mononucleotide compositions. Conversely, CpG dinucleotides were suppressed in vertebrates (*Chordata*) and molluscs (*Mollusca*) (**Fig 3A and B).**CpG suppression was even more evident in vertebrate mRNA 3’UTRs (**Fig 3C and S2 Fig A**), while the presence of CpGs in 5’UTRs was variable, more so than any other dinucleotide, perhaps reflecting the comparatively short length of many 5’UTRs and a role for RNA structure in translation regulation (**Fig 3C**). Overall, these data indicate that CpG-suppression is observed primarily in vertebrates, is not specific to coding sequences, and was likely present in animals prior to the emergence of genes closely related to the human ZAP CpG-specific antiviral gene.

**Fig 3.**
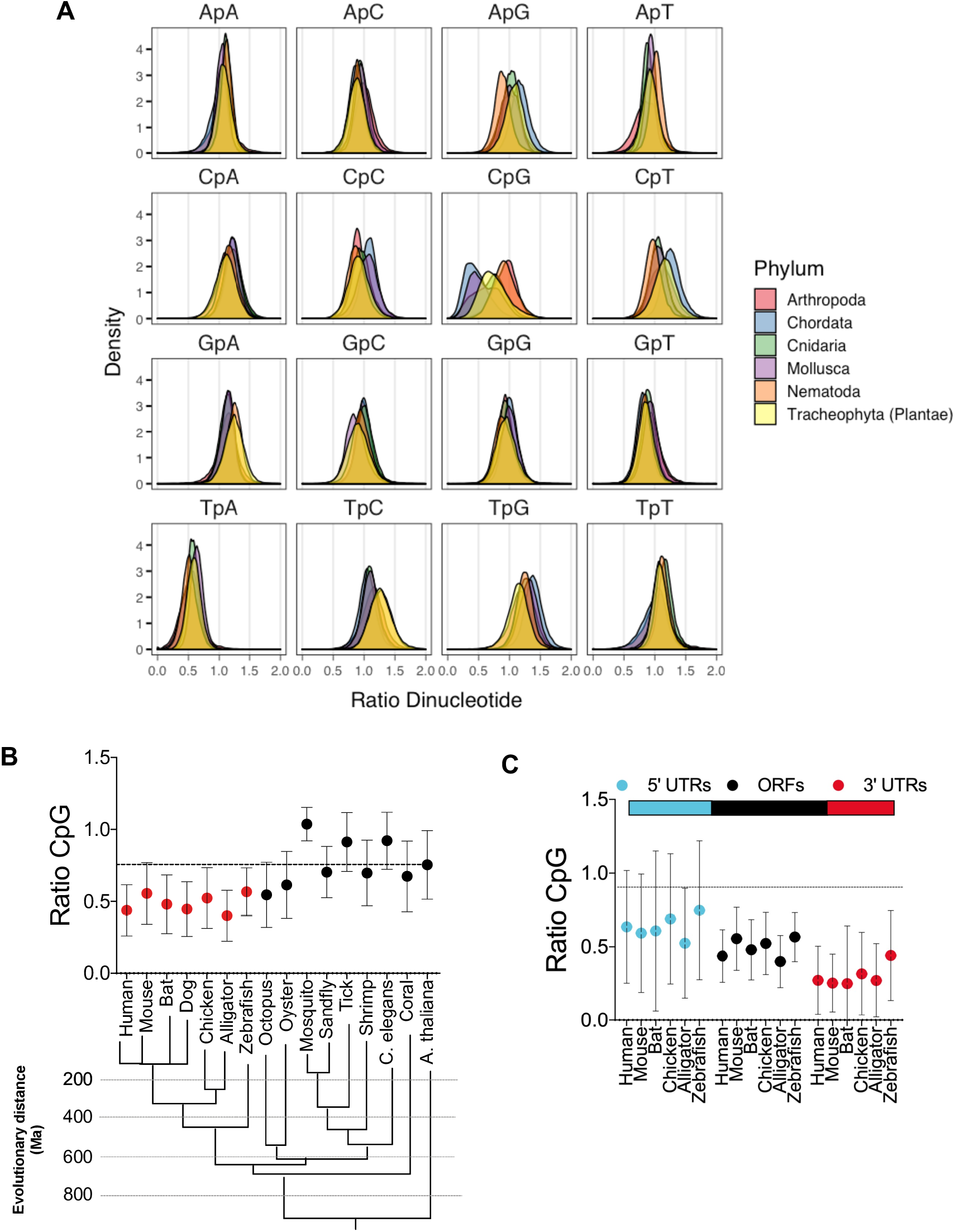
RNA transcripts from vertebrates are CpG-suppressed. (A) Open reading frames (ORFs) from several organisms belonging to the indicated phyla were collected from the NCBI nucleotide database and dinucleotide frequency ratio (observed/expected) was calculated and frequency distribution for each dinucleotide in ORFs was plotted. (B) Average dinucleotide observed/expected ratio in various animal species. Evolutionary distance (presented in million years, Ma) of indicated species was based on (34). (C) Average dinucleotide observed/expected ratio in various portions of vertebrate mRNA transcripts vertebrates.

### ZAP proteins from tetrapods have antiviral activity, but some require a cognate TRIM25

We next assessed whether the ZAP-related genes from tetrapods that share the highest homology with human ZAP exhibit CpG-targeted antiviral activity. During initial experiments, we found that the full-length ZAP proteins from various species were expressed at inconsistent levels in human cells. Therefore, in an attempt to circumvent this confounding variable, we constructed plasmids expressing chimeric proteins in which the N-terminal zinc finger domains of mouse, bat, chicken and alligator ZAPs were fused to the remaining portions (including the WWE and PARP-like domains) of human ZAP (**Fig 4A**). Importantly, the isolated N-terminal domain of ZAP has been reported to be sufficient for antiviral activity (3) and is responsible for RNA recognition. By constructing these chimeras we aimed to maintain the RNA-binding specificity of ZAP proteins while also maintaining the putative regulatory functions of the WWE and PARP-like domains of human ZAP. To test the activity of these chimeric proteins, we first transfected HEK293T ZAP^−/−^ cells with proviral plasmids and increasing amounts of plasmids encoding the various ZAP chimeras. Chimeric ZAP proteins with N-terminal domains (NTDs) from human (huZAP) and bat (baZAP) had antiviral activity against HIV-1_CG_, while ZAP chimeras with NTD’s derived from alligator (alZAP) and mouse (moZAP) were less potent, the ZAP chimera with an NTD from chicken (chZAP) had little or no antiviral activity (**Fig 4B and C**).

**Fig 4.**
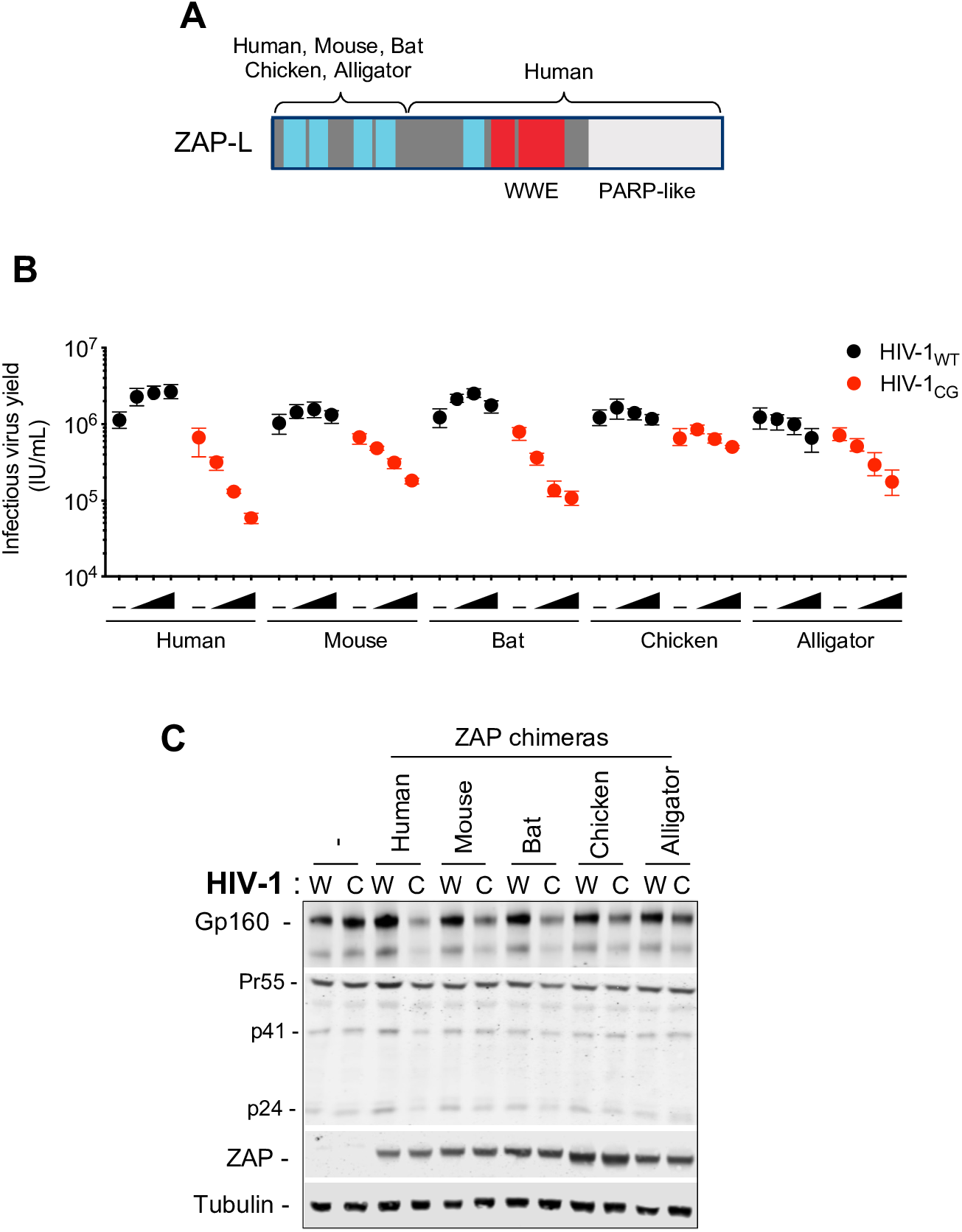
ZAP protein antiviral activity in human cells. (A) Schematic representation of ZAP chimeras in which the RNA binding domain of mouse, bat, chicken and alligator ZAP was fused to the WWE and PARP-like domain of human ZAP. (B-C) HEK293T ZAP^−/−^ cells were transfected with HIV-1_WT_ (W) or HIV-1_CG_ (C) proviral plasmids and increasing amounts (0ng, 75ng, 145ng, or 290ng) of a plasmid encoding each ZAP chimera. Supernatant was harvested after 48h and the infectious virus yield was determined using MT4-R5-GFP target cells (B). Whole cell lysates were analysed by western blotting probing with antibodies against HIV-1 proteins and ZAP (C). -, Indicates co-transfection with an empty vector in place of a ZAP expression plasmid.

The failure of the non-mammalian ZAP NTD chimeras to function in human cells could have been due to an absence of RNA recognition activity, or incompatibility with a required cofactor in human cells. One important co-factor for ZAP is TRIM25 (6,7). Even though HEK293T ZAP^−/−^ cells express TRIM25, the nature of the TRIM25-ZAP interaction and specifically, the ZAP domain with which TRIM25 functionally interacts is unknown.

Therefore, we explored whether the ZAP NTD interacts with TRIM25. Experiments in which ZAP-L and TRIM25 were overexpressed in HEK293T ZAP^−/−^ TRIM25^−/−^ cells (**Fig 5A**) revealed that ZAP and TRIM25, could be co-immunoprecipitated, as previously reported (6). Removing the TRIM25 SPRY domain (one truncated form was composed of the N-terminal 371 amino acids and the other was composed of the N-terminal 410 amino acids) dramatically reduced the HIV-1_CG_ -specific antiviral activity of ZAP (**Fig 5B**), as well as the ability of TRIM25 to co-immunoprecipitate with ZAP, again consistent with prior reports (**Fig 5A,B**) (6). Next we conducted coimmunoprecipitation experiments in which the ability of full-length human ZAP-L, a truncated form of ZAP lacking the NTD (ΔZnF ZAP-L) or a truncated form comprised only of the NTD (ZAP 254) to coimmunoprecipitate with TRIM25 was compared (**Fig 5C**). While full length ZAP and ZAP 254 coimmunoprecipitated with TRIM25, the inactive (**Fig 5D**) truncated ZAP, lacking the NTD, failed to coimmunoprecipitate with TRIM25 (**Fig 5C**). This data suggests that ZAP and TRIM25 functionally interact via the N-terminal ZAP zinc finger domain and the C-terminal SPRY domain of TRIM25.

**Fig 5.**
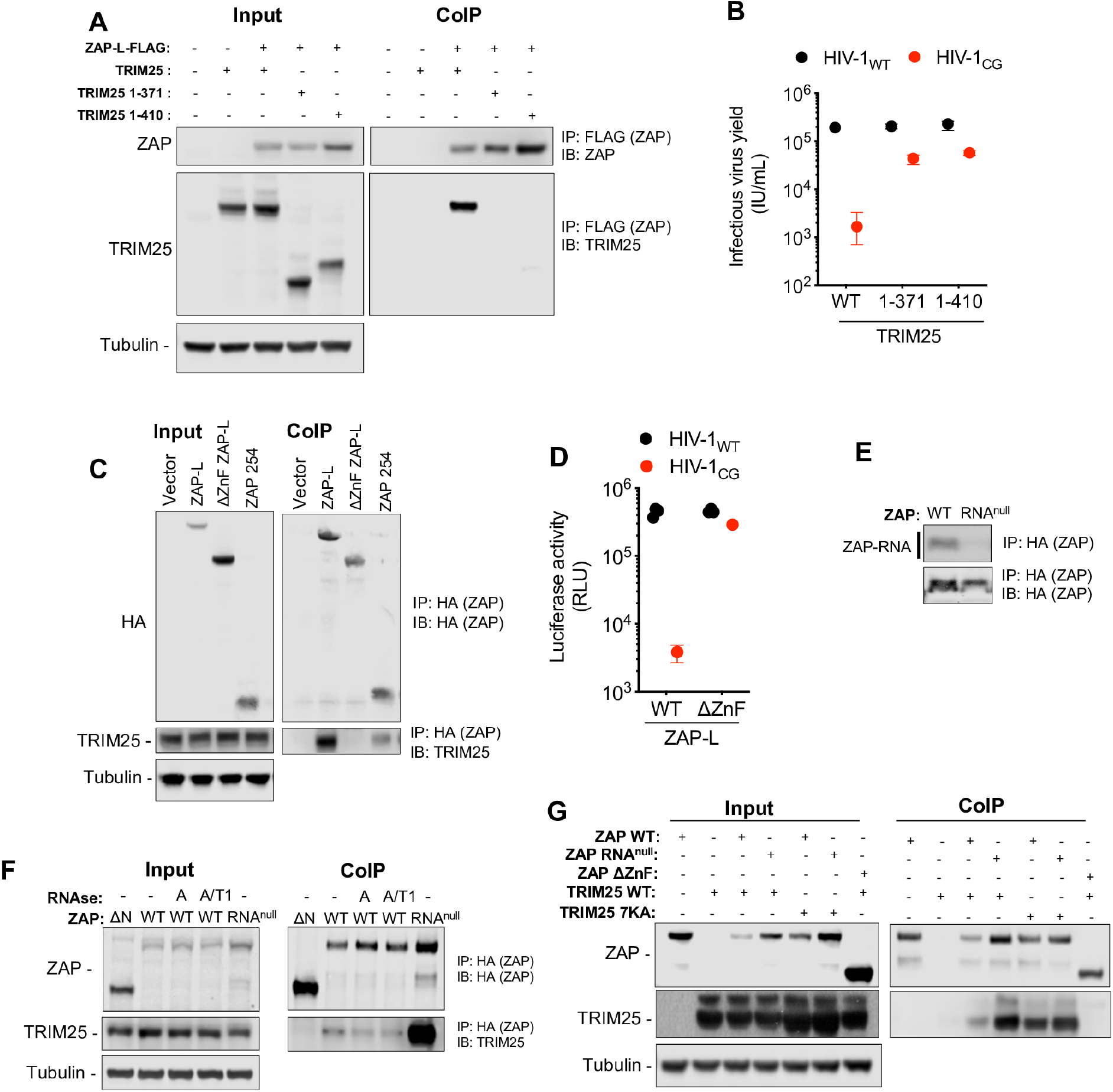
TRIM25 binds the ZAP NTD independently of RNA. (A) HEK293T ZAP^−/−^ and TRIM25^−/−^ cells were co-transfected with plasmids expressing human ZAP-L-FLAG and full-length untagged human TRIM25 or one of two human TRIM25 truncations (corresponding to the N-terminal 371 or 410 amino acids of TRIM25), that lack the SPRY domain (20). Proteins were immunoprecipitated from cell lysates with an anti-FLAG antibody and subjected to western blot analysis. IP, immunoprecipitation. IB, immunoblot. (B) HEK293T ZAP^−/−^ and TRIM25 ^−/−^ cells were co-transfected with proviral plasmids and plasmids encoding human ZAP and full-length TRIM25 or the indicated truncated TRIM25 proteins. Infectious virus yield was determined using MT4-R5-GFP target cells. (C) HEK293T ZAP^−/−^ cells were transfected with an empty vector or plasmids encoding human ZAP-L-HA, a truncated ZAP lacking the RNA binding domain (ΔZnF ZAP-L) or a truncated ZAP comprising the N-terminal 254 amino acids (ZAP 254). Proteins were immunoprecipitated from cell lysates with an anti-HA antibody and analysed by western blotting to detect overexpressed ZAP-L-HA and endogenous TRIM25. (D) Antiviral activity of ΔZnF ZAP-L was assessed by co-transfecting HEK293T ZAP^−/−^ cells with indicated proviral plasmids and plasmids encoding full length (WT) or truncated (ΔZnF) human ZAP. Infectious virus yield was determined using MT4-R5-GFP target cells. (E) HEK293T ZAP^−/−^ cells were transfected with plasmids encoding human ZAP-L (WT) or an RNA-binding mutant of ZAP (RNA^null^, R74A, R75A, K76A). After overnight incubation cells were irradiated with UV light, and ZAP proteins were immunoprecipitated. RNA bound to each protein was radiolabelled and protein-RNA adducts were resolved by SDS-PAGE, transferred to a nitrocellulose membrane and exposed to autoradiographic film. In parallel, a western blot of immunoprecipitated proteins was done using anti-HA antibody. IP, immunoprecipitation. IB, immunoblot. (F) HEK293T ZAP^−/−^ cells were transfected with plasmids encoding HA-tagged human ZAP-L (WT), ΔZnF ZAP-L or ZAP (RNA^null^). Cell lysates were treated with RNAse A or a mixture of RNAse A and T1. ZAP protein complexes immunoprecipitated and subjected to western blot analysis to detect overexpressed ZAP-L-HA and endogenous TRIM25. (G) HEK293T ZAP^−/−^ and TRIM25 ^−/−^ cells were transfected with plasmids encoding HA-tagged ZAP-L (WT), ΔZnF ZAP-L or ZAP (RNA^null^), along with untaggedTRIM25 or a TRIM25 RNA binding mutant (TRIM25 7KA). ZAP-HA protein complexes immunoprecipitated and subjected to western blot analysis to detect overexpressed ZAP and TRIM25 proteins.

Since both the ZAP NTD and TRIM25 proteins can bind RNA (11,20,21), it was possible that RNA might have a bridging function and be responsible for the interaction between the two proteins. To explore this hypothesis, we generated a ZAP mutant (R74A, R75A, K76A, hereafter referred to as ZAP RNA^null^). Notably ZAP RNA^null^ bound undetectable levels of RNA in crosslinking immunoprecipitation assays (**Fig 5E**). We transfected HEK293T ZAP^−/−^ cells with plasmids expressing wildtype ZAP-L or the ZAP RNA^null^ mutant, lysed the cells and treated the cell lysates with RNase A or a mixture of RNase A and T1 prior immunoprecipitation (**Fig 5F**). RNase A or A/T1 treatment only slightly affected coimmunoprecipitation of ZAP with endogenous TRIM25. Furthermore, the ZAP RNA^null^ mutation increased, rather than decreased, the amounts of coprecipitated endogenous TRIM25. Similarly, in experiments where HEK293T ZAP^−/−^ TRIM25^−/−^ cells were cotransfected with plasmids expressing wildtype or RNA-binding-defective mutants of ZAP (ZAP WT and ZAP RNA^null^) and TRIM25 (TRIM25 WT and TRIM25 7KA (20)), the amount of coprecipitated TRIM25 did not suggest a role for RNA in mediating ZAP-TRIM25 interactions (**Fig 5G**). Specifically, the amount of TRIM25 coprecipitated with ZAP was the same or increased when either ZAP RNA^null^ or TRIM25 7KA mutants were expressed in place of the wild-type proteins. Overall, these results suggest that ZAP NTD and TRIM25 interact in a manner that is dependent on protein-protein rather than protein-RNA interactions.

Since human TRIM25 interacted with human ZAP in an NTD dependent manner, but the ZAP chimeras encoded an NTD from a different species, it was possible that an inability to functionally interact with endogenous human TRIM25 was responsible for their absent or poor activity in of some ZAP chimeras in human cells. To explore this hypothesis, we tested whether the expression of TRIM25 from various species could support the activity of ZAP chimeras encoding a ZAP NTD from that species. TRIM25-related genes are present in mammals, birds and reptiles and some species of fish (**Fig 6A**). To assess whether the expression of cognate TRIM25 could augment the antiviral activity of each ZAP chimera, we measured infectious HIV-1_CG_ yield from HEK293T ZAP^−/−^ TRIM25^−/−^ cells cotransfected with plasmids expressing TRIM25 and chimeric ZAP proteins from various species (**Fig 6B**). While there was some variation in potency, ZAP chimeras from mammalian species generally exhibited antiviral activity in the presence of TRIM25, regardless of the species of origin. Conversely ZAP proteins from more distantly related species (chicken and alligator) were poorly active when coexpressed with mammalian TRIM25 proteins, but exhibited greater antiviral activity when coexpressed with a TRIM25 protein from the same species (**Fig 6B**).

**Fig 6.**
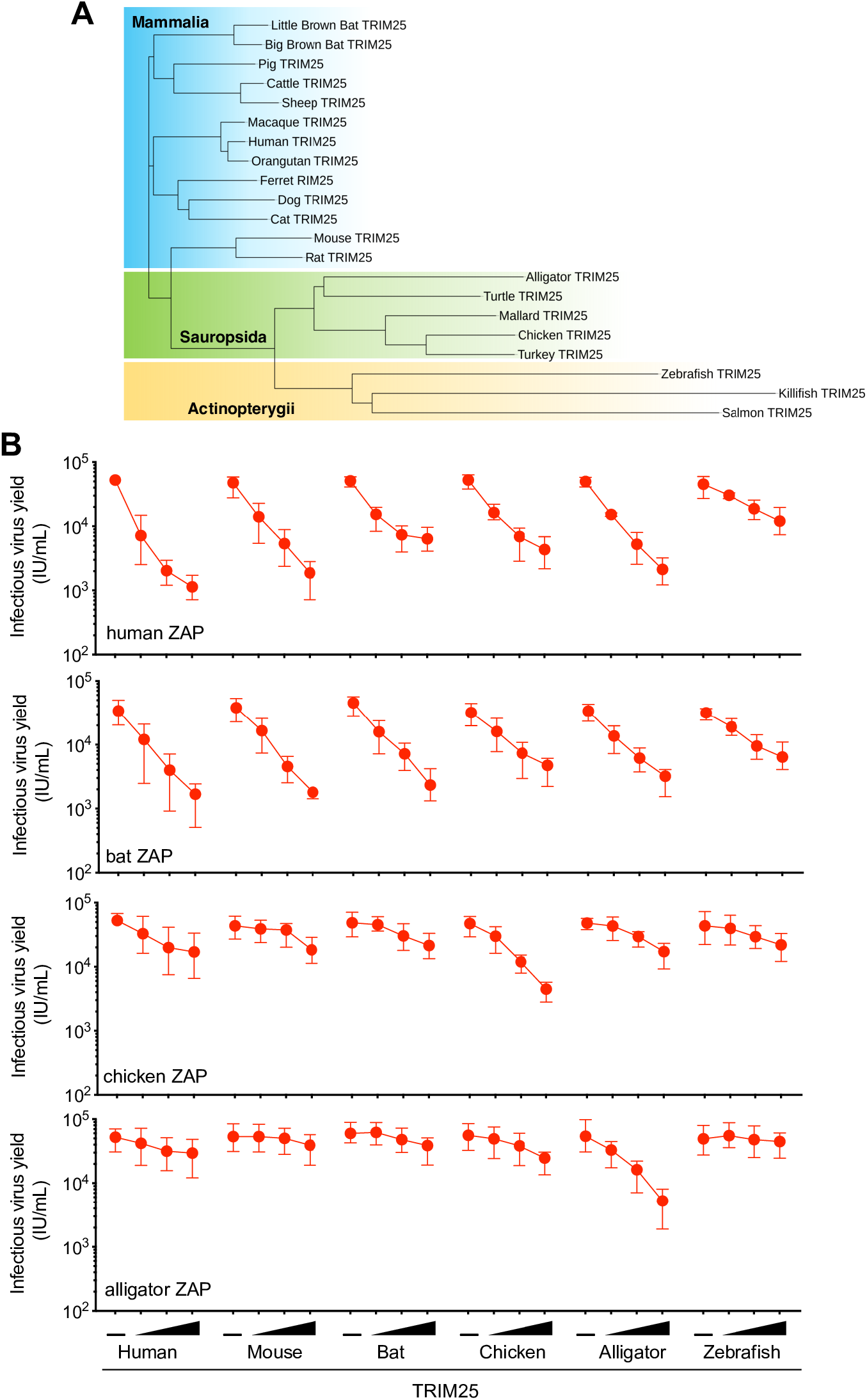
ZAP proteins from tetrapods have antiviral activity in the presence of cognate TRIM25. (A) Phylogenetic analysis of TRIM25 protein sequences from vertebrate species. Shaded areas indicate cluster sequences from mammals (in blue), birds and reptiles (in green) and ray-finned fish (in yellow). (B) HEK293T ZAP^−/−^ and TRIM25^−/−^ cells were co-transfected with an HIV-1_CG_ proviral plasmid as well as a fixed amount of a plasmid encoding human ZAP or ZAP chimeras. For each ZAP protein, cells were also co-transfected increasing amounts of a plasmid (0ng 10ng, 30ng or 90ng) expressing a TRIM25 protein from the 6 different indicated species. After 48h, infectious virus yield was determined using MT4-R5-GFP target cells.

To test whether the ZAP NTD from the various species had the same or different target RNA specificities, HEK293T ZAP^−/−^ and TRIM25^−/−^ were co-transfected with plasmids expressing a ZAP chimera and a cognate TRIM25 protein, together with a HIV-1_WT_ or HIV-1_CG_ proviral plasmid (**Fig 7A and B**). Interestingly, while all the ZAP/TRIM25 proteins reduced the yield of HIV-1_CG_, some of the TRIM25 and ZAP chimera combinations also reduced the yield of HIV-1_WT_ to some extent. This was particularly the case for the chicken TRIM25 and ZAP chimera combination which reduced the yield of infectious HIV-1_WT_ by >10 fold at the highest dose tested, suggesting that ZAP from chicken might have a different or expanded RNA binding specificity.

**Fig 7.**
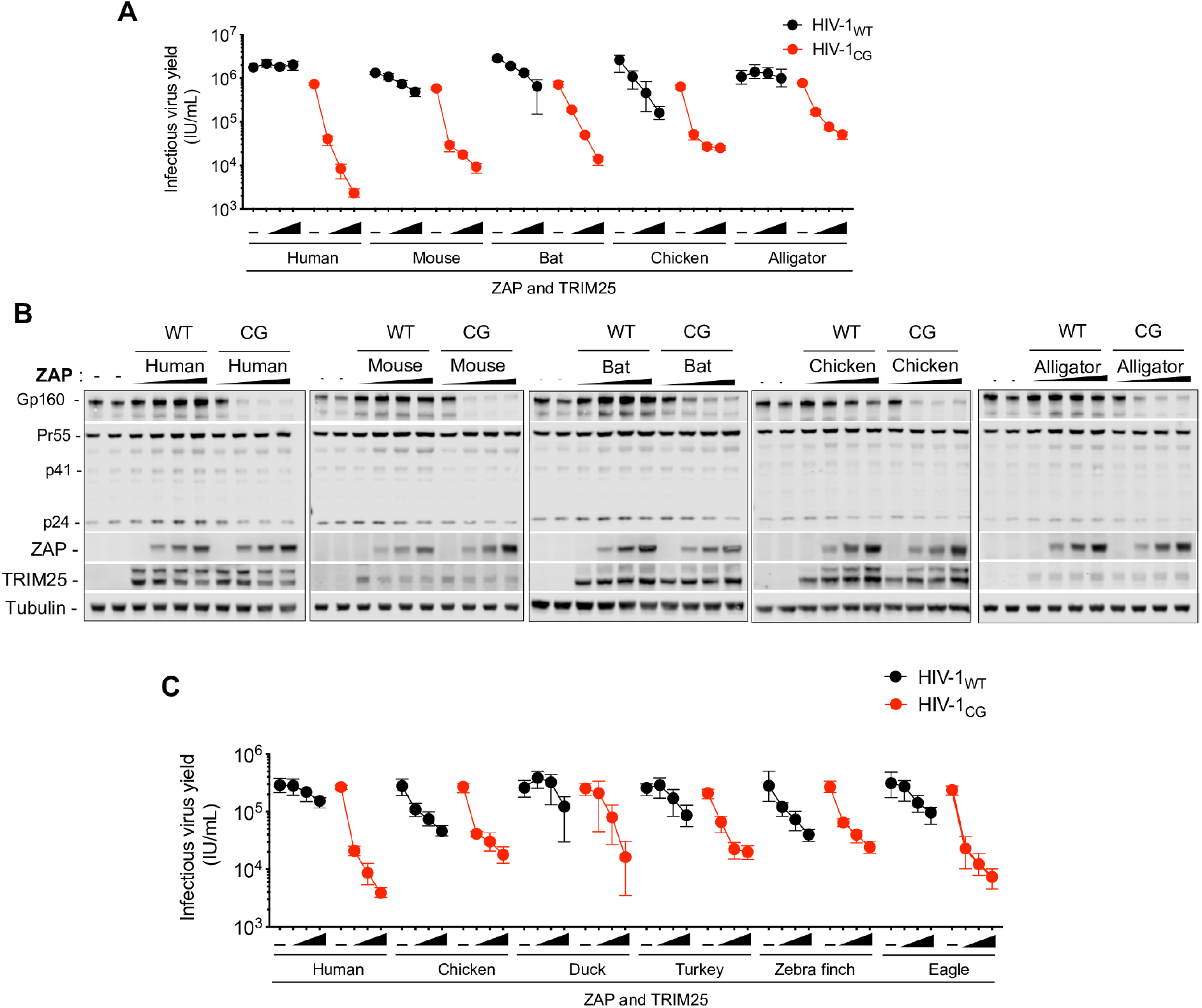
Avian ZAP proteins are less selective for CpG enriched HIV-1. (A-B) HEK293T ZAP^−/−^ and TRIM25^−/−^ cells were co-transfected with indicated proviral plasmids, increasing amounts (0ng, 75ng, 145ng, or 290ng) of a plasmid encoding an FLAG-tagged human ZAP or mouse, bat, chicken and alligator ZAP chimeras and a fixed amount of a plasmid encoding a cognate HA-TRIM25 protein. After 48h, infectious virus yield was determined using MT4-R5-GFP target cells (A). Whole cell lysates were analysed by western blotting probing with antibodies against HIV-1 proteins ZAP-FLAG and TRIM25-HA (B). (C) HE293T ZAP^−/−^ and TRIM25^−/−^ cells were transfected with HIV-1 proviral plasmids, along with increasing amounts (0ng, 75ng, 145ng, or 290ng) of a plasmids expressing human ZAP or ZAP chimeras from various avian species. For cells transfected with human ZAP, a fixed amount of a human TRIM25 expression plasmid was co-transfected, while for the avian ZAP chimeras, chicken TRIM25 was used. After 48h, infectious virus yield was determined using MT4-R5-GFP target cells.

Birds comprise one of the most diverse phyla in vertebrates, with about 10,000 species, adapted to a wide range of habitats (22). Since chicken ZAP was phenotypically distinct from other vertebrate ZAPs, we investigated whether ZAP from other species of birds share a similar phenotype with chicken ZAP. We constructed chimeric ZAP proteins in which the RNA binding domain was from one of several additional bird species: three from the order *Galliformes* – chicken (*Gallus gallus*), turkey (*Meleagris gallopavo*) and duck (*Anas platyrhynchos*) – and two from the suborder *Neoaves* – eagle (*Aquila chrysaetos canadensis*) and zebra finch (*Taeniopygia guttata*) (**Fig 7C and D**). The avian ZAP chimeric proteins exhibited different levels antiviral activity, but some, particularly the chicken and zebra finch ZAP chimeras reduced HIV-1_WT_ yield (**Fig 7B and D**). Indeed, the zebra finch ZAP chimera was nearly as potent against HIV-1_WT_ as it was against HIV-1_CG_. These findings suggest that a loss of selectivity for CG-rich RNA, or expansion of RNA target specificity has occurred on more than one occasion in avian ZAP proteins.

### RNA-binding specificity of chicken and human ZAP proteins differs

Since human ZAP binds to CG-rich RNA with high selectivity (10,11), differences in RNA binding affinity and specificity might explain the altered antiviral specificity of some avian ZAP proteins. We focused on chicken ZAP, and used crosslinking immunoprecipitation coupled with RNA sequencing (CLIP-Seq) to determine its RNA-binding preference in HEK293T ZAP^−/−^ TRIM25^−/−^ cells cotransfected with an HIV-1_CG_ proviral plasmid along with plasmids encoding human or chicken ZAP. This analysis revealed that, as expected and previously reported (10), human ZAP bound primarily to regions of the genome that contained a high number of CG dinucleotides (**Fig 8A**), Notably, the chicken chimeric ZAP preferentially bound to the CG-rich portion of the viral genome, but also exhibited a comparatively higher level of binding to the CpG-poor portions of the viral genome. We quantified the representation of each of the 16 dinucleotides in ZAP-crosslinked total reads from the viral genome for both human and chicken ZAP (**Fig 8B**). While CpG dinucleotides were enriched in RNAs crosslinked to both chicken and human ZAP chimeras, CpG enrichment was clearly less pronounced in chicken ZAP. Together these data suggest that chicken ZAP is less selective toward CG-rich RNA and this property corelates with the broader antiviral activity of chicken ZAP.

**Fig 8.**
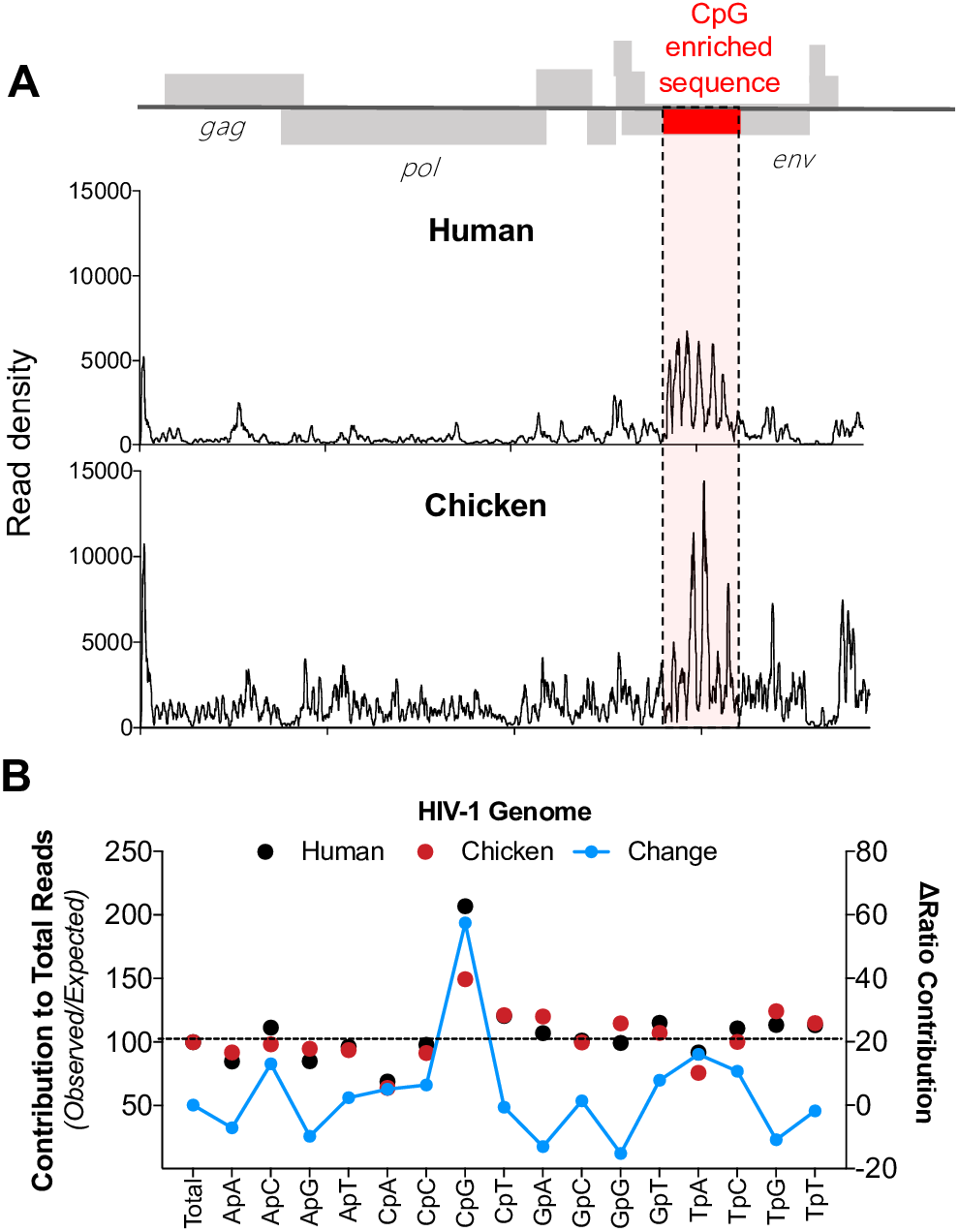
Differences in the RNA binding profiles of human and chicken ZAP. A) HEK293T ZAP^−/−^ and TRIM25^−/−^ cells were transfected with an HIV-1_CG_ proviral plasmid and a plasmid expressing human ZAP or chicken ZAP chimera. Cells were treated with 4-thiouiridine prior to UV-crosslinking and CLIP analysis. Reads associated with human (top) or chicken (bottom) ZAP were mapped to the HIV-1_CG_ genome and the read density plotted against position in the the HIV-1_CG_ genome. CpG-enriched region the HIV-1_CG_ genome is highlighted in red and dashed lines. (B) Contribution (% observed/expected) for each dinucleotide to reads derived from the HIV-1_CG_ genome to was calculated. Shift in the ratio for each dinucleotide contribution comparing human and chicken ZAP reads was also determined (blue line).

### A determinant in the ZAP NTD that governs species-specific TRIM25 dependence

The maximal antiviral activity of the chicken ZAP NTD chimera requires chicken TRIM25. In an attempt to determine what region in the ZAP NTD is responsible for this species-specific dependence, we inspected aligned protein sequences from human and chicken ZAP NTDs. We noticed that a divergent protein sequence, composed of two predicted α-helices is present in the C-terminal portion of the NTD (**Fig 9A, B**). Based on the crystal structure of human ZAP (11), these α-helices are located proximal to the third and fourth zinc fingers, facing away from the RNA-binding pocket (**Fig 9A**). To test whether this element is important for species-specific TRIM25 dependence, we generated a chimera that contained the four N-terminal zinc fingers from chicken ZAP, and the divergent NTD α-helices from the human ZAP (chZAP-X, **Fig 9B**). We compared the activity of chZAP-X with that of the human ZAP, and the previously constructed chZAP chimera in the presence of human or chicken TRIM25 (**Fig 9C**). As previously observed, human ZAP was active against HIV-1_CG_ in the presence of either human or chicken TRIM25. Conversely, the original chZAP chimera was more potent in the presence of chicken TRIM25 than human TRIM25, and also exhibited activity against HIV-1_WT_ in the presence of chicken TRIM25 (**Fig 9C**). Notably, the chZAP-X chimera, was more active against HIV-1_CG_ than chZAP in the presence of human TRIM25. Moreover, chZAP-X was also active against HIV-1_WT_ in the presence of either human or chicken TRIM25. Overall these data indicated that the four NTD zinc fingers of chicken ZAP are responsible for the its expanded antiviral activity against HIV-1_WT_, while the two divergent NTD α-helices are at least partly responsible for the species-specific dependence of chicken ZAP protein on chicken TRIM25.

**Fig 9.**
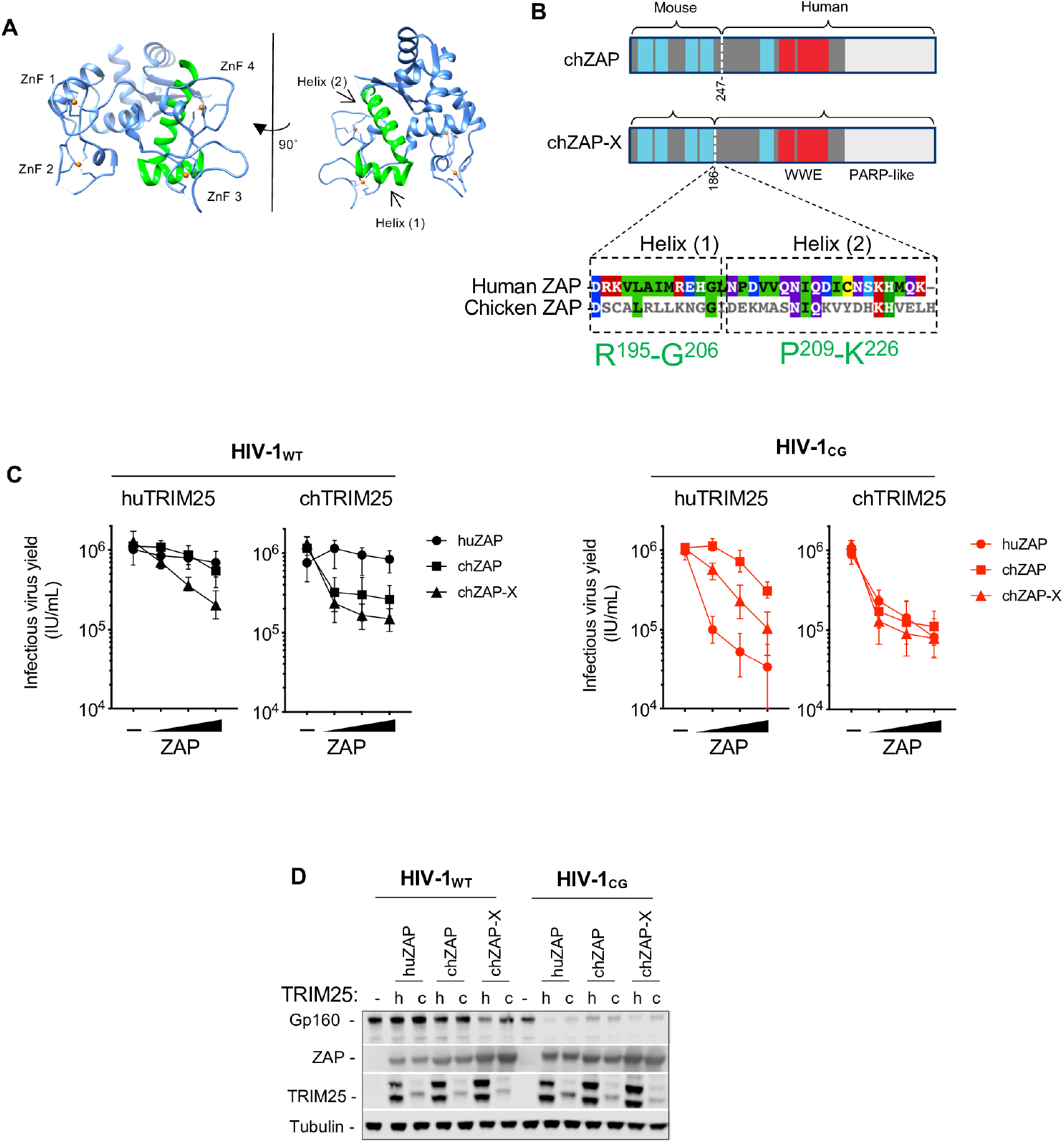
A determinant in the ZAP NTD contributes to the species-specific cognate TRIM25 requirement. (A) Representation of the crystal structure of the RNA-binding NTD domain of human ZAP (PDB 6UEI, (11)). ZnF 1-4 indicate zinc fingers 1 through 4. Areas colored in green indicate α-helices 1 and 2 at the C-terminus of the NTD. (B, top) Schematic diagram of chicken ZAP (chZAP) chimera, containing the N-terminal 247 amino acids of chicken ZAP, in an otherwise human ZAP background, and chicken ZAP-X (chZAP-X) chimera, that contains the N-terminal 186 amino acids of chicken ZAP in a human ZAP background. chZAP-X contains the two human α-helices highlighted (A). (B, bottom) Sequence alignment of the α-helices 1 and 2 in human ZAP with the corresponding region in chicken ZAP. Colors indicate amino acid identity and conservation. The position numbers for the two α-helices in humanZAP are indicated in green. (C-D) HEK293T ZAP^−/−^ and TRIM25^−/−^ cells were co-transfected with HIV-1_WT_ and HIV-1_CG_ proviral plasmids, as well as plasmids encoding human ZAP (huZAP), chicken ZAP chimera (chZAP) and the chicken ZAP-X (chZAP-X) chimera and either human TRIM25 or chicken TRIM25. After 48h, infectious virus yield was determined using MT4-R5-GFP target cells (C). Whole cell lysates were analysed by western blotting probing with antibodies against HIV-1 proteins ZAP-FLAG and TRIM25-HA (D)

## DISCUSSION

The emergence of genes with new functions is central to the adaptability of organisms to new challenges. Viral infections impose a strong selective pressure on their hosts, thus gene products with antiviral functions constitute prominent examples of genetic innovation and are among the fastest evolving genes (23). While our analysis suggests that *ZAP* originated from a duplication of the *PARP12* gene, and both human PARP12 and ZAP have been reported to exert antiviral activity against a broad range of viruses, the mechanisms by which they exert their function are different. Specifically, ZAP antiviral function is proteins is primarily related to their RNA recognition (3,4,10), while PARP12 showed no detectable RNA-binding activity. Indeed previous work has indicated that PARP12 antiviral activity is linked to PARP domain-dependent ADP-ribosylation of viral proteins (16,24). The PARP-like domain of mammalian ZAP proteins is catalytically inactive, and therefore, cannot catalyze ADP-ribosylation of viral or endogenous proteins (25,26). Interestingly, however, sequence alignments of PARP domains of chicken and alligator ZAP, revealed that these ZAP proteins likely contain catalytically active PARPs, raising the possibility that these are bifunctional antiviral proteins. A more recent gene duplication has apparently occurred in mammals, as revealed by the existence of another paralogue of ZAP, termed ZAP-like or *ZC3HAV1L*. This gene encodes a shorter protein, that contains four CCCH-type zinc fingers and appears paralogous to the ZAP RNA-binding domain (**S1 Fig**). Since the ZAP RNA-binding domain is sufficient to inhibit virus replication (3), it is plausible that ZC3HAV1L has antiviral activity. However, this awaits definitive experimentation.

Our prior work suggests that ZAP exploits the CpG suppressed state of human genomes to discriminate between self and non-self RNA (10). CpG suppression in animal genomes is thought to be the by-product of DNA methylation and subsequent deamination of 5-methyl-cytosine into thymidine, followed by the repair of the mispaired G on the opposing strand (13). Thus, the extent to which CpG dinucleotides in a given genome is methylated is linked to the degree of CpG suppression therein. Indeed, while the dinucleotide frequency distributions are similar for most dinucleotides in most organisms, CpG dinucleotide frequency in animal genomes can vary widely (**Fig 3A**). In particular, the coding sequences of several vertebrates (mammals, birds, reptiles, fish) and some non-arthropod invertebrates (molluscs) show substantial levels of CpG suppression, while in arthropods, nematodes and some species of plants the frequency of CpG dinucleotides is higher. This property correlates with the extent of DNA methylation observed in these organisms (27). Thus, the CpG suppressed state of animal genomes creates the opportunity for self-nonself discrimination by ZAP-related proteins that has apparently been exploited in tetrapods. Nevertheless, one study showed the rate of CpG DNA methylation in genomes may be insufficient to explain the extent to which vertebrate mRNA is CpG-suppressed (28). Indeed, RNA transcripts in some organisms seem to be under additional selective pressures to purge CpG dinucleotides, as suggested by the observed frequency of C-to-A mutations in a CpG context. Thus, it is possible that the emergence of ZAP/PARP12 related genes may have further shaped the dinucleotide composition of coding sequences in modern genomes.

We found that the activity of ZAP proteins from different organisms was inherently dependent on the cellular context. Indeed, in line with previous reports (6,7) we found that the expression of a ZAP cofactor, TRIM25, was important for its antiviral activity against CpG-enriched HIV-1. In fact, the activity of ZAP proteins from certain species (chicken and alligator) was dependent on the coexpression of a cognate TRIM25 protein. This species-specific dependency suggests that ZAP and TRIM25 interact in a way that is maintained across different species. Concordantly, replacing two poorly conserved, α-helices immediately C-terminal to the RNA binding domain in chicken ZAP with the equivalent element from human ZAP enabled the chimeric protein to function better in the presence of human TRIM25. This result strongly suggests a species-specific component of the interaction between ZAP and TRIM25. Consistent with this finding, prior work, confirmed herein, has suggested a physical interaction between ZAP and TRIM25 in a manner that was dependent on the TRIM25 SPRY domain (6). Additionally, we showed that the ZAP NTD, that includes the RNA binding domain, is necessary and sufficient to co-immunoprecipitate TRIM25. Notably, two domains in TRIM25 have been reported to bind RNA: (1) a short section of positively-charged amino acids located between the coiled-coil domain and the PRY/SPRY domain (N-KKVSKEEKKSKK-C, amino acids 381-391 (20)), and (2) a small region (470-508) within the PRY/SPRY domain (21)). Additionally, two reports suggested that RNA enhances the E3-ubiquitin ligase activity of TRIM25 *in vitro* (20,21). However, we found that a TRIM25 mutant in which the lysine cluster at 381-391 was replaced by alanines (TRIM25 7KA) supported antiviral activity. Moreover, TRIM25 7KA, ZAP-RNA^Null^ and RNAseA/T1 treatment were all compatible with ZAP-TRIM25 coprecipitation. Together, these results suggest that ZAP-TRIM25 interaction is not RNA-dependent and is likely mediated by protein-protein contacts. That there exists a species-specific restriction in ZAP-TRIM25 functional compatibility, that can be mapped to a specific protein element, is consistent with this notion.

Finally, we found that avian ZAP proteins, unlike human ZAP, exhibit varying degrees of antiviral activity against HIV-1_WT_ as well as HIV-1_CG_. Consistent with this finding, CLIP-Seq analysis using chicken ZAP revealed that it binds more promiscuously throughout the viral genome than does human ZAP. Recently, two structures of ZAP (from human and mouse) bound to a target RNA elucidated the nature of the interaction between ZAP and CG dinucleotides (11,12). Key contacts are established by ZAP residues K89, Y98, K107 and Y108 (11). Of these, K89, K107 and Y108 are conserved between human and chicken ZAP. Notably, the second zinc finger of chicken ZAP is three-amino acids shorter than the human counterpart; how the structural divergence between these human and avian ZAP influences RNA binding specificity required further investigation, but mutations at various positions surrounding the CpG binding pocket in human ZAP can cause relaxation in strict CpG specificity (11). Interestingly, birds are important vectors for certain human viral infections, such as influenza and West Nile virus. The CpG dinucleotide content of influenza A viruses has progressively decreased in human populations following zoonotic transmission from birds (29). Moreover, the frequency of CG dinucleotides in viral genomes can be used to predict animal reservoirs (30). In the case of birds, viruses that infect *Neoaves* – and to some extent viruses that infect *Galliformes* – have higher CpG dinucleotide frequencies than viruses that infect primates or rodents. Together, our results provide a potential molecular explanation for the observed fluctuations in CpG-content of viruses that have adapted to different hosts, and suggest that ZAP’s antiviral activity represents a selective pressure that influences the dinucleotide composition of viral genomes.

Overall, these findings highlight the potential for innovation of gene function driven by viral infection. Understanding how different organisms have evolved to control infections will illuminate the mechanisms of host range restriction and enable novel interventions in viral diseases.

## Materials and Methods

### Cells

Human embryonic kidney (HEK) 293T, HEK293T ZAP^−/−^ (10), HEK293T ZAP^−/−^ TRIM25^−/−^ (6) and chicken fibroblasts DF1 (ATCC, CRL-12203) were cultured in Dulbecco’s Modified Eagle’s Medium (DMEM) supplemented with foetal bovine serum (FBS). *Eptesicus fuscus* kidney (EFK) cells were purchased from Kerafast, Inc (ESA001) and cultured in DMEM supplemented with FBS. Mouse embryonic fibroblasts (MEFs, (31)) were cultured in DMEM supplemented with bovine calf serum (BCS). MT4-R5-GFP cells (32) were grown in RPMI medium supplemented with FBS. All cells were maintained at 37°C in 5% CO_2_.

### Plasmids

Sequences encoding the RNA-binding domains of alligator, turkey, duck, zebra finch and eagle ZAP were retrieved from the NCBI nucleotide database and DNA was synthesised by Twist Biosciences. Total RNA from EFK cells was extracted using the NucleoSpin RNA purification kit (Macherey-Nagel) according to manufacturer’s guidelines, reverse transcribed into cDNA using the SuperScript III First-Strand Synthesis System (Invitrogen), and bat ZAP was amplified using specific primers. TRIM25 mRNA was isolated from MEF, EFK, DF1 cell lines, as above, and reverse transcribed to generate cDNA. Alligator, turkey, duck, zebra finch and eagle TRIM25 cDNA was synthesised by Twist Biosciences. Total RNA from the intestine of a zebrafish specimen (kindly provided by Professor A. James Hudspeth, The Rockefeller University) was extracted and TRIM25 cDNA was generated as before. ZAP chimeras were generated by fusing the RNA-binding domain of ZAP isolated from indicated species to the residues 255-902 of the long isoform of human ZAP-L, that was C-terminally tagged with three HA epitopes or one FLAG epitope, as indicated, and inserted into the expression plasmid pCR3.1. TRIM25 sequences were fused to a C-terminal HA epitope and inserted into pCR3.1. The two truncated versions of TRIM25 (1-371 and 1-410) and the TRIM25 7KA mutants were a kind gift from Owen Pornillos, University of Virginia (20). The ZAP RNA^null^ mutant was generated by introducing three point-mutations R74A, R75A and K76A. by overlap extension PCR

### Bioinformatics

The protein sequences of human ZAP and PARP12 were retrieved from GeneBank. These sequences were subsequently used to identify orthologues in other species using the Blastp suite of NCBI. Sequences with significant E-values were used for subsequent sequence alignments and phylogenetic analysis. We considered the identified protein sequences to be products of orthologues of *ZC3HAV1*/*PARP12* if (1) significant sequence homology was observed, (2) if sequences contained 5 CCCH-type zinc fingers and (3) if WWE and PARP-like domains were present. MUSCLE was used to perform multiple sequence analysis. Phylogenetic trees were derived using BEAST (33) and plotted using FigTree. Speciation dates were based on (34). A similar approach was taken to identify and analyse *TRIM25* orthologues. For synteny studies, *ZC3HAV1/PARP12* loci from different vertebrate species were analysed using Ensembl and Genomicus Browsers.

For nucleotide composition studies, RNA transcript sequences from human (*Homo sapiens*), house mouse (*Mus musculus*), little brown bat (*Myotis lucifugus*), chicken (*Gallus gallus*), alligator (*Alligator mississippiensis*), dog (*Canis familiaris*), zebrafish (*Danio rerio*), California two-spot octopus (*Octopus bimaculoides*), oyster *(Crassostrea gigas),* mosquito (*Aedes aegypti*), sandfly (*Lutzomyia longipalpis*), tick (*Ixodes scapularis*), pacific white shrimp (*Litopenaeus vannamei*), *Caenorhabditis elegans*, coral (*Acropora digitifera*) and *Arabidopsis thaliana* were retrieved from the NCBI nucleotide sequence database. Subsequently, sequences were fragmented into 5’ UTRs, ORFs and 3’UTRs. Mononucleotide and dinucleotide frequencies were calculated using the SSE suite (35) and analysed using in-house R scripts.

### Virus Yield Assays

To assess the activity of different zinc finger proteins, virus yield was measured as described previously(11). In brief, HEK293T ZAP^−/−^ TRIM25^−/−^ were seeded onto a 24-well plate and transfected with 375ng of HIV-1_WT_ or HIV-1_CG_ proviral plasmids, along with 87ng of a plasmid encoding TRIM25 and varying amount of ZAP expression constructs, unless otherwise indicated. After 24h, media were replaced and at 48h post-transfection supernatants were collected, filtered through a 0.22μm filter and titres determined using MT4-R5-GFP target cells. Transfected cells were lysed in NuPAGE buffer and protein samples were analysed by western blotting.

### Luciferase Assays

Expression constructs encoding a recoded, low-CpG firefly luciferase and containing wildtype or CG-enriched VSV-G or IAV-NP sequences as 3’ UTRs were used to assess the activity of ZAP and PARP12, as described before (10). Cells were co-transfected with 50ng of luciferase-encoding plasmids, 250ng of plasmids encoding human ZAP-L or PARP12 and human TRIM25. After 56h, cells were lysed in cell lysis buffer and luciferase activity was measured using the Luciferase Assay System (Promega).

### Western Blotting

Cell lysates were incubated at 72°C for 20min and sonicated for 20s. Protein samples were resolved in a 4-12% PAGE gel (Novex) using MOPS running buffer. Protein was transferred to a nitrocellulose membrane, blocked at room temperature and incubated with the following antibodies: anti-ZC3HAV1 (rabbit, 1:10,000, clone 16820-1-AP, Proteintech Group), anti-TRIM25, anti-HA (rabbit, 1:5000, clone 600-401-384, Rockland), anti-Tubulin (mouse, 1:10,000, clone DM1A T9026, Millipore-Sigma), anti-HIV-1 Env (goat, 1:1000, 12-6205-1, American Research Products). Blots were washed and incubated with secondary antibodies: anti-Mouse IgG IR700 Dye Conjugated (Licor), Anti-Rabbit IgG IR800 Dye conjugated (Licor), Anti-Goat IgG IR800 Dye Conjugated (Licor) and Anti-Rabbit IgG horseradish peroxidase conjugated (Jackson). Blots were imaged immediately in a Licor Odyssey scanner or incubated with ECL substrate and imaged on a CDigit blot scanner.

### RNA-Protein Immunoprecipitation

Cells were seeded onto a 15cm dish and transfected, 24h later, with 8μg of plasmids encoding ZAP-L-3xHA, PARP12-3xHA or ZAP RNA^Null^ (R74A,R75A, K76A)-3xHA. The following day, media were replaced by culture media containing 4-thiouridine. Two days after transfection, cells were exposed to UVB light (0.15 J cm^−2,^ λ = 365 nm, Stratalinker 2400 UV), washed in PBS and lysed in Lysis Buffer (10mM HEPES pH7.5, 30mM KCl, 40μM EDTA, 0.1% Igepal CA-630 supplemented with protease inhibitor cocktails). Lysates were clarified by centrifugation and incubated with RNAse A for 5min at 37°C. Anti-HA mouse antibody (BioLegend) was adsorbed to protein G agarose beads and incubated with the cell lysates for 2h at 4°C. Beads were washed twice in NP40 Lysis Buffer, twice IP wash buffer (50mM HEPES pH 7.5, 300mM KCl, 2mM EDTA, 0.5% Igepal CA-630), twice in LiCl buffer (250mM LiCl, 10mM Tris pH 8.0, 1mM EDTA, 0.5% Igepal CA-630, 0.5% sodium deoxycolate), twice in NaCl Buffer (50mM Tris pH 7.5, 1M NaCL, 1mM EDTA, 0.1 SDS, 0.5% sodium deoxycholate) and twice in KCl buffer (50mM HEPES pH 7.5, 500 mM KCl, 0.05% Igepal CA-630). RNA:Protein complexes were incubated with calf intestinal phosphatase (NEB) for 13min at 37°C and washed in phosphatase wash buffer (50mM Tris HCl pH 7.5, 20mM EGTA, 0.5% NP40). Beads were resuspended in PNK buffer (50 mM Tris HCl pH 7.5, 50mM NaCl, 10mM MgCl_2_) and incubated with 5 U of PNK in the presence of 0.5 μCi/μL γ-^32^P ATP. Beads were washed, lysed in NuPAGE Lysis buffer and RNA:Protein complexes were resolved in a 4-12% NuPAGE gel. Complexes were transferred to nitrocellulose membrane and exposed to autoradiographic film.

### Co-immunoprecipitation

To evaluate ZAP and TRIM25 interactions, cells (HEK293T ZAP^−/−^ or HEK293T ZAP^−/−^ TRIM25^−/−^) were seeded onto a 10cm dish and co-transfected, 24h later, with 3μg of a plasmid encoding ZAP-FLAG and 3μg of a plasmid encoding TRIM25-3xHA. In experiments in which endogenous TRIM25-positive cells were transfected, only ZAP-encoding plasmids were used. Cells were lysed 48h after transfection in 1.5mL of Lysis Buffer (10mM HEPES pH7.5, 30mM KCl, 40μM EDTA, 0.1% Igepal CA-630 supplemented with protease inhibitor cocktails). Lysates were clarified by centrifugation and treated with RNAse A (Roche, 100 Units) or RNAse A/T1 (NEB, 100 Units) for 5min at 37°C. Protein G agorose beads that were pre-adsorbed to either anti-HA (BioLegend) or anti-FLAG (Millipore) antibodies were added to the lysates and incubated at 4°C for 2h. Magnetic beads were captured and washed twice in lysis buffer and three times in IP wash buffer (50mM HEPES pH 7.5, 300mM KCl, 2mM EDTA, 0.5% Igepal CA-630). Protein complexes were resuspended in NuPAGE buffer and resolved on a 4-12% PAGE gel, transferred to nitrocellulose membranes and analysed by western blotting.

### CLIP-Seq

RNA:protein complexes were isolated as described above. RNA was isolated and prepared for sequencing as before (36). In brief, after isolation of RNA:protein complexes, RNA was released using Proteinase K (Roche). Purified RNA fragments were ligated to 3’ and 5’ adaptors, reverse transcribed (SuperScript First-Strand cDNA Synthesis System, Invitrogen) and amplified by PCR. The resulting cDNA library was sequenced using the Illumina HiSeq 2000 platform. Sequencing reads were processed and analysed as described previously (10).

---

**Fig S1.**
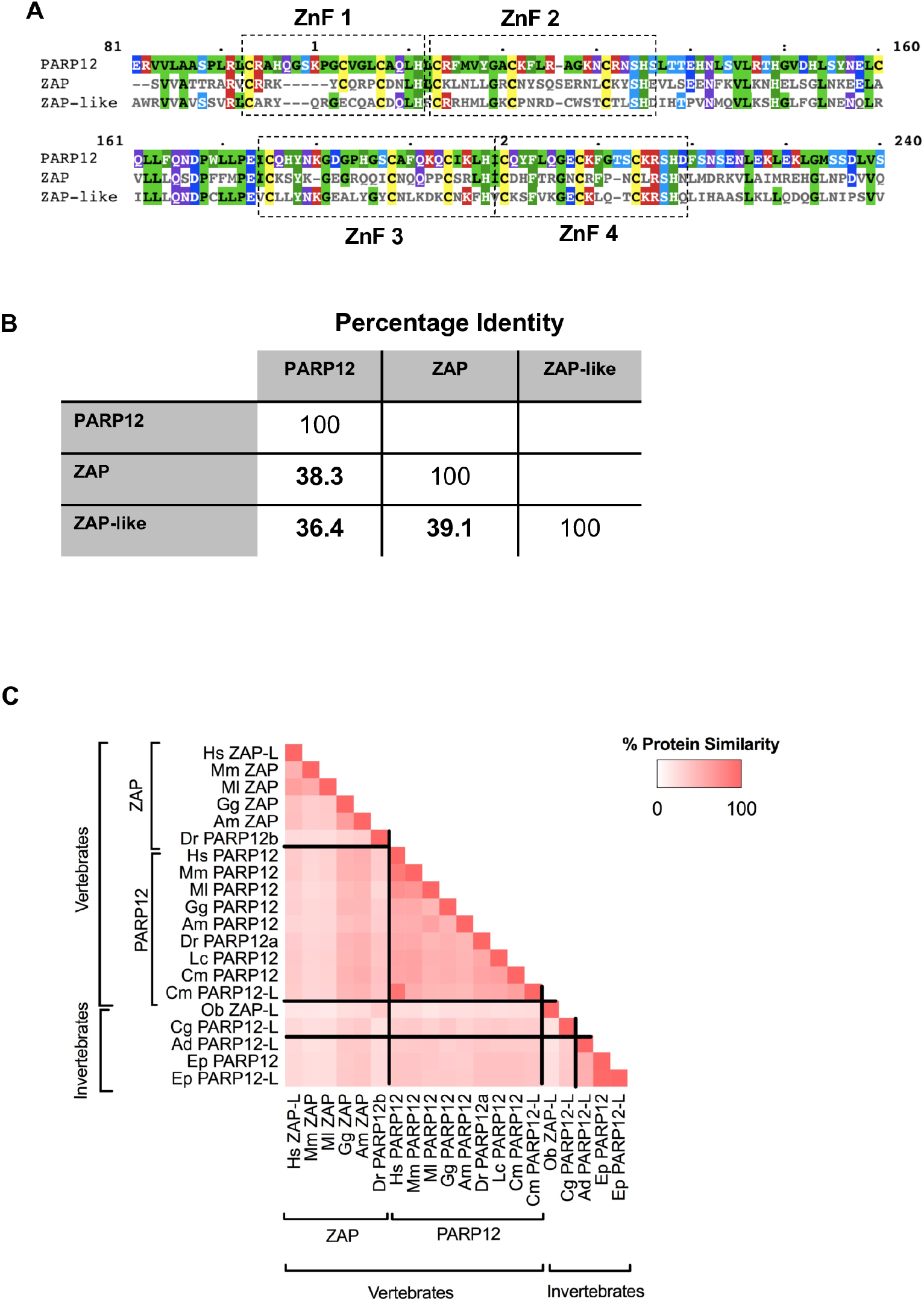
Homology among ZAP, PARP12 and ZAP-like proteins. (A) Protein sequence alignment of the N-terminal domain (NTDs) of human ZAP, PARP12 and ZAP-like protein. Colored residues indicate amino acid properties and conservation. (B) Percentage identity matrix among human ZAP, PARP12 and ZAP-like proteins. (C) Percentage protein similarity one-to-one matrix among ZAP and PARP12 paralogues found in vertebrate and invertebrates species. Hs, *Homo sapiens* (human); Mm, *Mus musculus* (mouse); Ml, *Myotis lucifugus* (little brown bat), Gg, *Gallus gallus* (chicken); Am, *Alligator mississipiensis* (alligator); Dr, *Danio rerio* (zebrafish); Lc, *Latimeria chalumnae* (Coelacanth); Ob, *Octopus bimaculoides* (California two-spot octopus); Cg, *Crassostrea gigas* (oyster); Ad, *Acropora digitifera* (coral).

**Fig S2.**
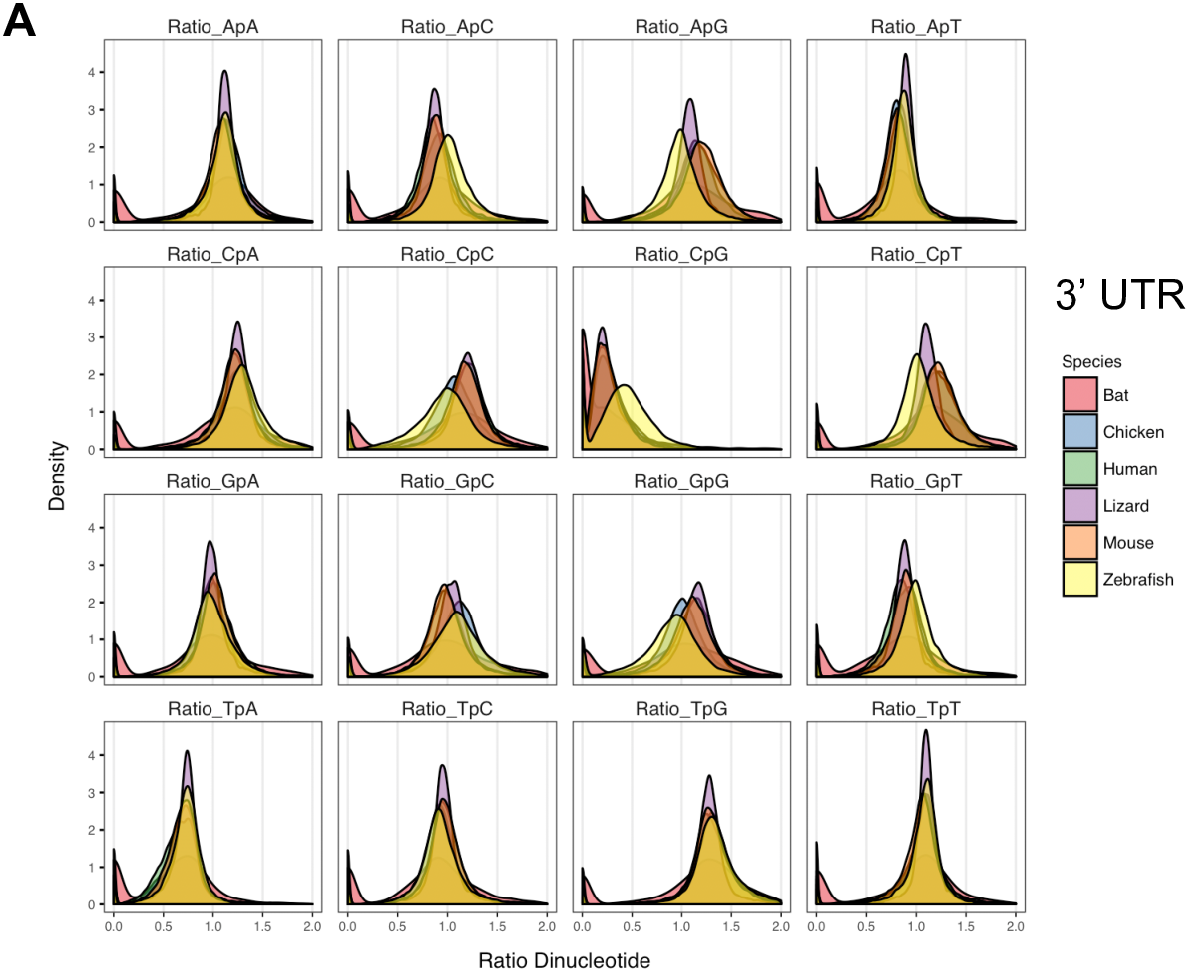
Dinucleotide composition in mRNA 3’UTRs across vertebrates. (A) The 3’ untranslated regions of mRNA transcripts found in transcriptomes of several vertebrates were collected from the NCBI nucleotide database and dinucleotide frequency ratio (observed/expected) was calculated and frequency distribution for each dinucleotide was plotted.

